# A rapid and flexible microneutralization assay for serological assessment of influenza viruses

**DOI:** 10.1101/2023.02.02.526820

**Authors:** Kalee E. Rumfelt, William J. Fitzsimmons, Rachel Truscon, Arnold S. Monto, Emily T. Martin, Adam S. Lauring

**Affiliations:** Department of Epidemiology, University of Michigan, Ann Arbor, MI, USA; Division of Infectious Diseases, Department of Internal Medicine, University of Michigan, Ann Arbor, MI, USA; Department of Microbiology and Immunology, University of Michigan, Ann Arbor, MI, USA

**Author notes:** **Corresponding Author:** Adam Lauring, MS2 4742C, SPC 1621, 1137 Catherine Street, Ann Arbor, MI 48109,.

**Keywords:** influenza virus, serology, neutralization, hemagglutination

## Abstract

**Background:** Serological responses from influenza vaccination or infection are typically measured by Hemagglutinin Inhibition (HAI) or Microneutralization (MN). Both methods are limited in feasibility, standardization, and generalizability. We describe a luciferase MN (LMN) assay that combines the advantages of the conventional MN assay with the ease of the HAI assay

**Methods:** Sera were obtained from the HIVE study. Plasmid transfection was used to generate recombinant influenza viruses expressing the hemagglutinin and neuraminidase of test strains, all other viral proteins from A/WSN33, and a NanoLuc reporter. Serum neutralization of luciferase-expressing targets was quantified as a reduction in Relative Light Unit (RLU) emission from infected cells. Neutralization titers were measured for cell- and egg-adapted versions of A/H3N2/Hong Kong/4801/2014 and A/H3N2/Singapore/INFIMH-16-0019/2016 and compared to HAI titers against egg-grown antigens.

**Results:** 333 sera were collected from 259 participants between May 2016 and July 2018. Sampled participants ranged from 7 to 68 years of age and >80% were vaccinated against influenza. HAI and LMN titers were correlated for A/H3N2/Hong Kong/4801/2014 (ρ=0.45, p≤0.01) and A/H3N2/Singapore/INFIMH-16-0019/2016 (ρ=0.75, p≤0.01). LMN titers were lower for cell strains compared to egg strains (A/H3N2/Hong Kong/4801/2014 mean log_2_ fold change = −2.66, p≤0.01 and A/H3N2/Singapore/INFIMH-16-0019/2016 mean log_2_ fold change= −3.15, p≤0.01).

**Conclusions:** The LMN assay was feasible using limited sample volumes and was able to differentiate small antigenic differences, specifically between egg-adapted and cell-derived strains. The correspondence of these results with the commonly-used HAI confirms the potential utility of this assay for high-throughput studies of correlates of protection and vaccine response.

## Introduction

Influenza causes 9 million to 36 million illnesses and 4,000 to 60,000 deaths in the United States (US) each year^1^. Vaccination is one of the best ways to prevent influenza infection and the large public health burden associated with it^2,3^. Influenza vaccination has been shown to have variable effectiveness from year to year, a phenomenon that has been attributed to egg-adaptations occurring during the manufacturing process that hamper the mounting of antibodies to circulating strains^4–9^. For this reason, it is critical to supplement observational studies investigating vaccine effectiveness, with serological analysis of vaccinated and unvaccinated individuals that are able to distinguish between antibodies elicited by egg versus cell-grown antigens.

Serological responses from influenza vaccination or infection can be investigated through multiple assays^10–12^. The Hemagglutinin Inhibition (HAI) assay measures the ability of antibodies to inhibit the binding of the virus’ Hemagglutinin (HA) surface glycoprotein to sialic acids on the surface of red blood cells (RBCs), thereby preventing hemagglutination^10^. The HAI assay is relatively simple and inexpensive, and it has been considered a gold standard of influenza serology for decades^10,11,13^. However, HAI assays are vulnerable to non-specific interfering factors, rely on availability of mammalian or avian fresh red blood cells, have a subjective readout method, and have limited utility for newer H3N2 strains^10–14^. More recently, microneutralization (MN) assays and focus reduction neutralization tests (FRNT) have become more common, as they are more sensitive than HAI assays and more broadly measure the neutralizing activity of sera^11,12,14^. The MN assay also performs well across more recent H3N2 strains^13^. Although MN assays have advantages over the HAI assay, they are more labor intensive and protocols are not well standardized across laboratories^11,12,14^.

Here we describe a luciferase-based MN (LMN) assay that combines the advantages of the conventional MN assay with the ease of the HAI assay. We generated recombinant influenza virus targets that express a luciferase reporter, providing a sensitive and quantitative read-out of cellular infection and the inhibitory activity of sera. We demonstrate this assay’s performance using cell culture- and egg-adapted versions of two recent H3N2 strains. We related the LMN assay’s detection of neutralization activity against egg-adapted H3N2 targets to the HAI assay detection of seroreactivity for the same targets and sera. The cell-adapted targets were used to compare the cell-adapted and the egg-adapted LMN neutralization responses. Together our results demonstrate the utility of this assay for studies of immune correlates of protection.

## Methods

### Participants and Sera

Sera were provided from individuals enrolled in the Household Influenza Vaccine Evaluation (HIVE) study. The HIVE study was approved by the Institutional Review Board at the University of Michigan Medical School (HUM00198212), and written informed consent was provided by patients or a proxy/surrogate. Methods pertaining to the HIVE study can be found elsewhere^15^. Briefly, individuals were followed and contacted weekly to identify acute respiratory infections (ARI). Blood draws were scheduled at enrollment and twice annually after that. Serum was separated via centrifugation. The sera were stored at minus 20°C until further processing. In total, we used 333 serum samples from 259 individuals between May 2016 and July 2018.

### Hemagglutinin Inhibition Assays

All serum samples underwent HAI testing as a standard processing method. The HAI assays were performed in 96-well plates by combining a standardized quantity of HA from specific A/H3N2, A/H1N1, and B vaccine strains with serially diluted sera and turkey red blood cells as previously described^15^. HA targets were obtained through the International Reagent Resource (IRR) and from Sanofi.

### Molecular cloning

Viral stocks for A/H3N2/Hong Kong/4801/2014 (FR-1453) and A/H3N2/Singapore/INFIMH-16-0019/2016 (FR-1590) strains were obtained from IRR. Stocks were passaged once at low multiplicity on MDCK-SIAT1 cells and RNA was extracted using the QIAmp Viral RNA mini kit (Qiagen). Genomic segments encoding the HA and NA proteins were amplified by reverse transcription polymerase (RT-PCR) with primers 5’-TATTCGTCTCAGGGAGCAAAAGCAGGGG-3’ and 5’-ATATCGTCTCGTATTAGTAGAAACAAGGGTGTTTT-3’ for HA and primers 5’-TATTGGTCTCAGGGAGCAAAAGCAGGAGT-3’ and 5’-ATATGGTCTCGTATTAGTAGAAACAAGGAGTTTTTT-3’ for NA. These primers contain BsmBI and BsaI restriction sites to facilitate cloning into pHW2000 (a gift from Robert Webster, St. Jude Children’s Research Hospital). The egg-adapted sequences of these plasmid clones were verified by Sanger sequencing. To create the cell-adapted strains, HA and NA gene synthesis was performed by Integrated DNA Technologies (IDT) for A/H3N2/Hong Kong/4801/2014. Only HA contained cell-adapted mutations for A/H3N2/Singapore/INFIMH-16-0019/2016, which was also synthesized by IDT. The synthesized segments contained BsmBI and BsaI restriction sites to facilitate cloning into pHW2000. The cell-adapted sequences of these plasmid clones were verified by Sanger Sequencing.

### Rescue of recombinant luciferase expressing influenza viruses

For transfections, 5×10^4^ HEK 293T and 2.5×10^4^ MDCK-SIAT1 cells were plated in 12 well plates in growth media (DMEM, Invitrogen #19965-092, with a final concentration of 10% Fetal Bovine Serum (FBS, Invitrogen #26140-079), 100 units/mL Penicillin (Invitrogen #15140-122), 100ug/mL Streptomycin (Invitrogen #15140-122), 2mM L-Glutamine (Invitrogen #25030-081)) the day prior to transfection^16^. These co-cultures were transfected with 500ng each of PASTN-PA-NanoLuc (a gift from Andrew Mehle, University of Wisconsin), strain specific pHW184-HA, strain specific pHW186-NA, pHW181-PB2, pHW182-PB1, pHW185-NP, pHW187-M, pHW188-NS and TransIT-LT1 (MIRUS #MIR2300). After 12 hours, the cultures were washed and seeded with viral media (DMEM, Invitrogen #19965-092, with a final concentration of 0.1875% Bovine Serum Albumin (BSA, Invitrogen #15260-037), 25mM HEPES (Invitrogen #15630-080), 100 units/mL Penicillin (Invitrogen #15140-122), 100ug/mL Streptomycin (Invitrogen #15140-122), 0.1 ug/mL TPCK-treated Trypsin (Worthington Biochemical #LS003740)). Once full cytopathic effect was observed, supernatants were centrifuged at 1400xg for 4 minutes and stored in a final concentration of 0.5% glycerol at −80°C. These passage 0 stocks were titered and used to infect 1×10^6^ MDCK-SIAT1 at an MOI of 0.01. These passage 1 supernatants were harvested as above and stored at −80°C in single use aliquots. Stocks were titered as median tissue culture infectious dose (TCID_50_) in 96-well plates, by scoring positive wells as luciferase expression that exceeded two times the cellular background. This titering method was consistent with conventional TCID_50_ measurements that scored cytopathic effect.

### Luciferase microneutralization assay

Human (positive) control serum was obtained from a vaccinated donor and stored at −20°C. Lyophilized sheep (negative) control serum was obtained from IRR, reconstituted using sterile water, and stored at −20°C. Before use, all serum aliquots were thawed at room temperature and resuspended by vortexing. Human control, sheep control, and patient sera were heat inactivated (HI) at 56°C for 30 minutes prior to use.

Control and patient sera were added to column 1 at a volume of 10μl and serially diluted across the plate in 100μl total volume (Figure 1A). For each target virus, the P1 stock was diluted to a concentration of 2×10^3^ TCID50/mL and 50μl (100 infectious units) were added to all the serum dilution wells. A back titer series was created by adding to 50μl (100 infectious units) to the first well in a column and then performing two-fold serial dilutions down to the last well. After a one-hour incubation at 37°C, 100μl MDCK-SIAT1 cells at a concentration of 1.5×10^5^ cells/mL were added to each well. The plate was then incubated for 18 hours at 37°C.

**Figure 1.**
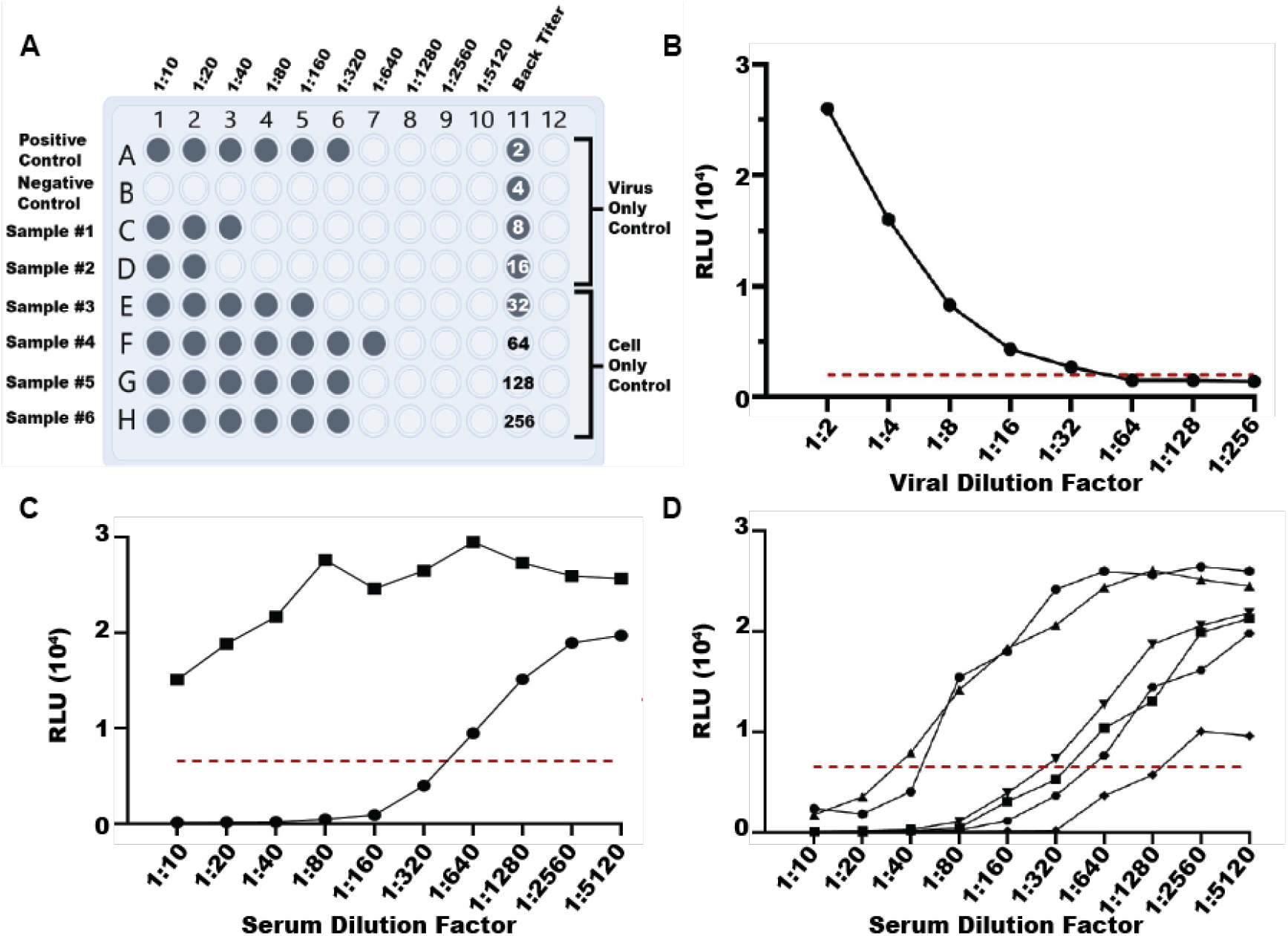
Design of the luciferase microneutralization assay (LMN). (A) Plate layout: Sera wells (rows A-H, columns 1-10) are considered positive for 75% neutralization of virus if the relative light unit (RLU) output is above the virus control RLU average minus the cell control RLU average divided by 4. Numbers in back titer wells indicate fold dilution of the input virus (100 TCID50). A back titer well is considered positive if the RLU output was greater than twice the cell only control RLU average. A plate was considered “passing” if the first negative well fell between the 5th and 8th dilution (1:32 to 1:256). Positive sera and back titer wells are colored. (B) Example of back titer with RLU (y-axis) for each virus dilution (x-axis). A visible leveling off in back titer between the 5^th^ and 8^th^ dilution was required for a plate to pass quality control. The red line denotes the cutoff of the cell only control average multiplied by two. (C) Example of positive (human) neutralization control (circles) and negative (sheep) neutralization control (squares). The 75% cutoff for neutralization is denoted by a red line. (D) Example of titrations for six sera, each denoted by a different shape. The 75% cutoff for neutralization is denoted by a red line.

After incubation, the supernatants were aspirated from the wells and 50μl of a 1:8000 concentration of ViviRen (Promega, #E6491) in viral media were added to each well. The plate was incubated at room temperature for 3 minutes, and luminescence was measured in a Synergy HTX multi-mode luminometer using a gain of 160. All the steps involving ViviRen were performed in a darkened room.

### Statistical analysis

Pearson Correlation Coefficients (PCC) were calculated to determine correlations between LMN and HAI titers separately for the A/H3N2/Hong Kong/4801/2014 and the A/H3N2/Singapore/INFIMH-16-0019/2016 strains. Paired t-tests were used to compare LMN neutralization titer against the egg-versus cell-adapted strains of A/H3N2/Hong Kong/4801/2014 and A/H3N2/Singapore/INFIMH-16-0019/2016. All statistical analyses were performed using SAS software version 9.4 (SAS Institute, Cary, NC).

## Results

The LMN assay is modeled after a traditional MN assay, which measures neutralization of virus in the presence of serum with potentially protective antibodies. In the LMN, influenza virus targets express the NanoLuc reporter from segment 3 (encoding PA). We used standard influenza virus reverse genetics to generate recombinant 6+2 viruses with segments expressing the hemagglutinin (HA) and neuraminidase (NA) of test strains and all other proteins, including the reporter, from A/WSN33 (H1N1). Expression of the NanoLuc reporter was sufficient to detect a single infectious event at 18 hours post-infection, as titers of viral stocks measured by luminescence over background matched titers measured by cytopathic effect (NanoLuc TCID50 vs. TCID50, data not shown).

As in the traditional MN assay, the LMN assay is performed in 96 well plates with 100 infectious units of target virus in each well and serial dilutions of test sera (Figure 1A).^11^ The plate has dedicated wells for “virus only controls (VC)” to capture maximum luminescent output, “cell only controls (CC)” to measure background, and two-fold serial dilutions of the target virus as a “back titer” to ensure that 100 infectious units were used. For a plate to pass quality control, the luminescent output (RLU) in the back titer wells should be less than two times the CC average in at least one well between wells 5 and 8 (viral dilutions of 1:32-1:256).

The 75% neutralization titer (NT_75_) is calculated as the serum dilution at which luminescence is reduced 75% relative to the VC (maximum luminescence in the absence of serum) and CC (background cellular luminescence) wells (Figure 1A). In preliminary testing, we found that the 75% cutoff was more discriminatory than a 50% cutoff given cellular background and occasional flat, as opposed to sigmoidal, curves with non-neutralizing control sera.

We evaluated the performance of the LMN assay using 333 serum specimens from 259 individuals collected before and after the 2016/2017 and/or the 2017/2018 Northern hemisphere influenza seasons. Thirty-seven individuals had paired pre- and post-season specimens for the same season, and some contributed 3 or 4 total specimens (Table 1). These sera were chosen, because they exhibited discordant HAI assay results for A/H3N2/Hong Kong/4801/2014 and A/H3N2/Singapore/INFIMH-16-0019/2016, two related but antigenically distinct, vaccine strains. The sampled individuals ranged from 7 to 68 years of age with many being between 41 and 64 years of age (49%). The sera came from a heavily vaccinated cohort with 81.4% and 86.5% of individuals having received the recommended annual influenza vaccine prior to the 2016-2017 and 2017-2018 influenza seasons, respectively.

**Table 1.**
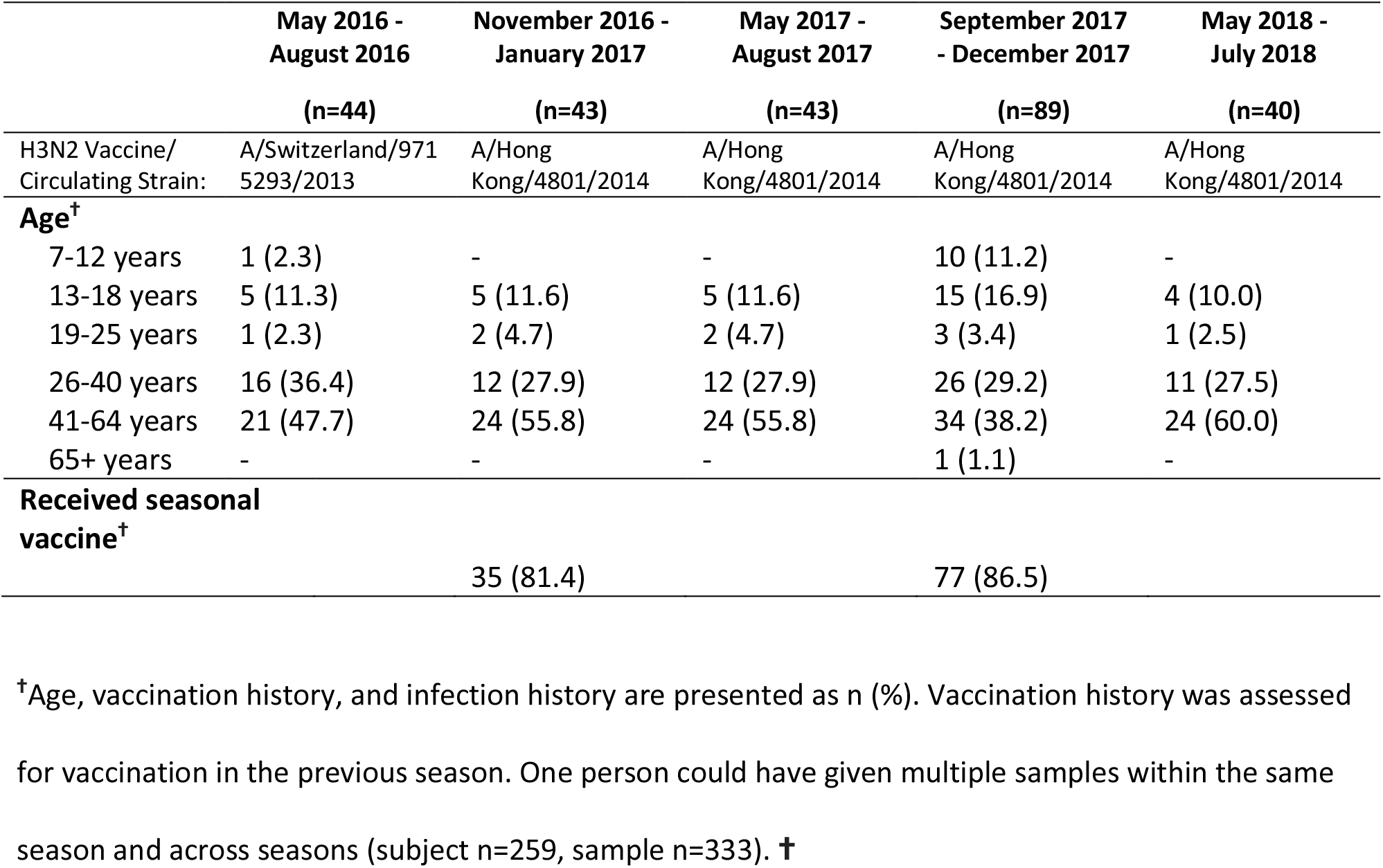
Demographic characteristics, vaccination history, and infection history of sampled individuals.

We compared LMN NT_75_ to HAI titer for each of these sera. Because these assays differ in their sensitivity, specificity, and dynamic range, both were ordinalized using a common approach. The NT_75_ for a negative control serum (e.g., sheep serum) was set to 0 and each two-fold serial dilution of the test sera was therefore 1, 2, 3, etc. For example, a LMN result with a negative control NT_75_ of 1:40 and a test serum NT_75_ of 1:160, would be expressed as 2. After ordinalization, we found that the HAI titer and the LMN NT_75_ were significantly correlated, but the magnitude differed by strain. Titers were moderately correlated for A/H3N2/Hong Kong/4801/2014 (PCC 0.45, p≤0.01; Figure 2A) and strongly correlated for A/H3N2/Singapore/INFIMH-16-0019/2016 (PCC 0.75, p≤0.01; Figure 2B).

**Figure 2.**
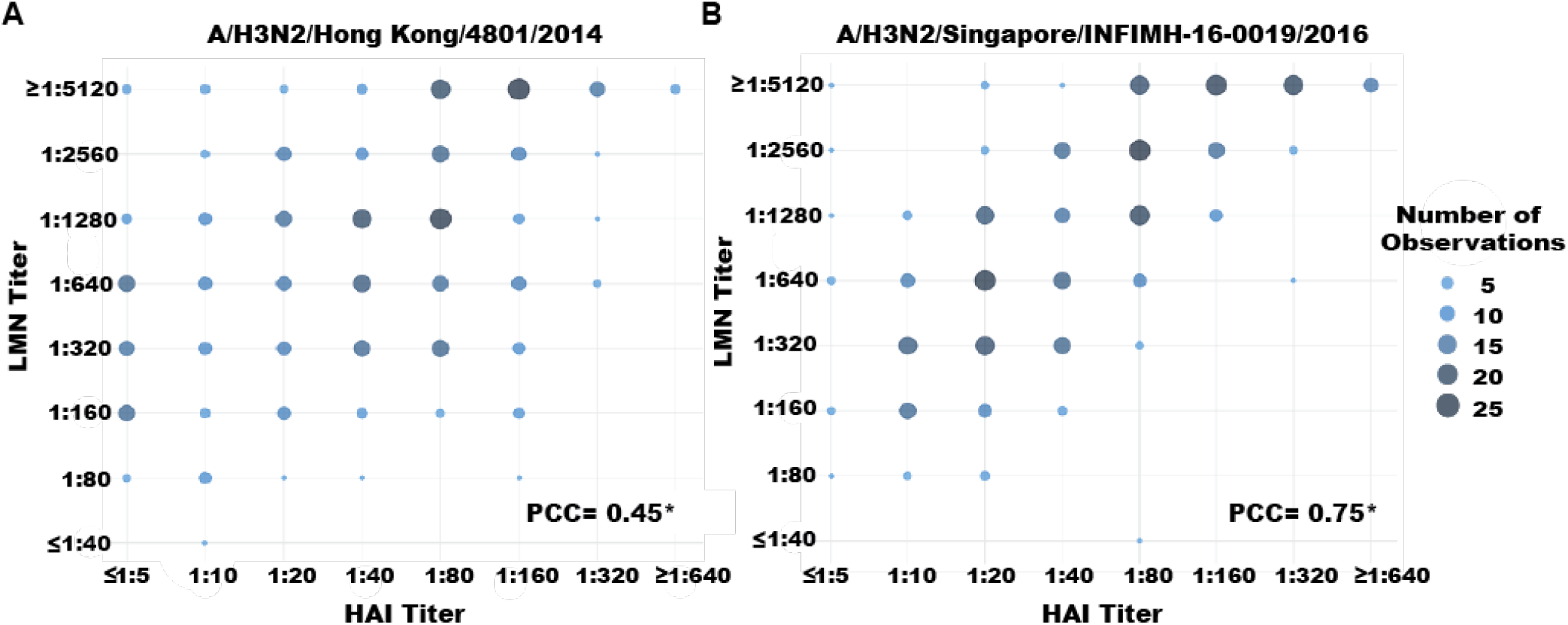
Correlation between HAI and LMN NT_75_ for (A) A/H3N2/HongKong/4801/2014 and (B) A/H3N2/Singapore/INFIMH-16-0019/2016; *Pearson Correlation Coefficients with a p-value ≤0.01.

We measured LMN NT_75_ against cell culture- and egg-adapted A/H3N2/Hong Kong/4801/2014 and A/H3N2/Singapore/INFIMH-16-0019/2016 influenza viruses to determine if the LMN assay could detect antigenic differences due to the corresponding mutations in HA and NA. We created egg-adapted versions of these strains by using HA and NA segments that corresponded to sequences of egg passaged viruses (see Methods). Because we rescued and propagated these egg-adapted strains in MDCK cells, we sequenced all stocks to ensure that the egg- and cell culture-associated mutations were maintained. The LMN detected differences in ordinalized LMN NT_75_ for cell culture- and egg-adapted strains of A/H3N2/Hong Kong/4801/2014 (mean log_2_ fold change= −2.66; Figures 3A and 3C) and A/H3N2/Singapore/INFIMH-16-0019/2016 (mean log_2_ fold change= −3.15; Figures 3B and 3D). All sera had higher NT_75_ for the egg-adapted strains except two sera against A/H3N2/Hong Kong/4801/2014 and one serum against A/H3N2/Singapore/INFIMH-16-0019/2016 (Figure 3).

**Figure 3.**
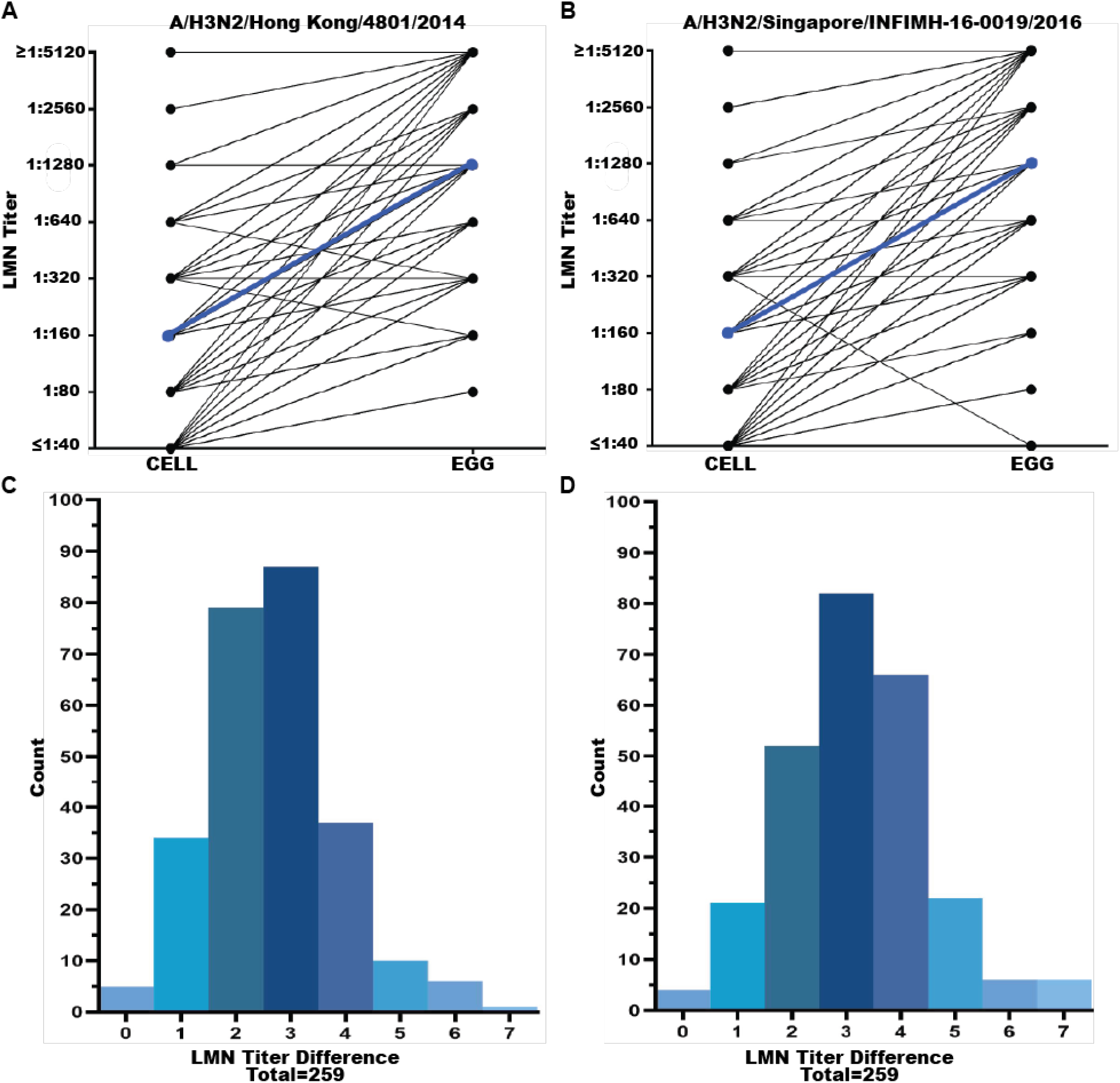
Difference in paired profile of LMN NT_75_ across egg- and cell-adapted influenza viruses. (A) and (B) Mean log_2_ fold change in LMN NT_75_ from cell- to egg-adapted viral targets is denoted by a blue line. Both the Hong Kong and Singapore strains showed a statistically significant different LMN NT_75_ for cell- and egg-adapted virus strains (Hong Kong t-value= −34.36, dF=258, p-value ≤0.01; Singapore t-value= −36.26, dF=258, p-value ≤0.01). (C) and (D) Mean log_2_ fold difference in LMN NT_75_ across cell- and egg-adapted virus strains (Hong Kong mean log_2_ fold change= −2.66 (−2.82, −2.51); Singapore mean log_2_ fold change= −3.15 (−3.33, −2.98)). All individuals had higher LMN NT_75_ for the egg-adapted strains except two individuals with a 1-fold difference for the A/H3N2/Hong Kong/4801/2014 and one individual with a 3-fold difference for A/H3N2/Singapore/INFIMH-16-0019/2016.

## Discussion

In this study, we present a new luciferase-based MN assay which performs similarly to the gold-standard HAI assay. We observed significant correlations between the LMN NT_75_ and HAI titer for two recent H3N2 strains, A/H3N2/Hong Kong/4801/2014 and A/H3N2/Singapore/INFIMH-16-0019/2016. Additionally, we were able to show that the LMN assay can detect differences in LMN NT_75_ for genetically similar egg-adapted and cell-adapted influenza viruses. Our results demonstrate that the LMN assay is a promising alternative to the HAI assay and is applicable to a broader range of research questions, including the study of subtle antigenic differences between strains that may have a major impact on seroprotection.

HAI titers are widely used as a serologic correlate of protection, although some have expressed concerns regarding their utility^12,13,17–19^. The LMN holds key advantages over the HAI assay. Specifically, the HAI assay has a subjective readout method, which can limit reproducibility, and it can produce variable results using recent H3N2 influenza strains^13^. The LMN assay addresses reproducibility issues by using the back titer, a built-in standardization component that provides a secondary check to ensure that the proper concentration of virus was added to each well on the plate^11^. This prevents extrapolating information from plates that were subject to experimental error. Additionally, the readout from the luminometer and calculation of the NT_75_ is performed objectively using pre-specified RLU cutoffs, eliminating subjective assessments of positive vs. negative wells. Another promising attribute of the LMN is that it performs well for many recent H3N2 strains (e.g. A/Hongkong/4801/2014), which do not hemagglutinate adequately in standard HAI assays^13^.

Although the LMN assay is similar to the traditional MN assay, it is easier to perform and quicker than the original. Generating target viruses for the LMN is relatively straightforward using standard influenza virus reverse genetics with cloned HA and NA genes, and targets can be produced in cells or eggs^20^. This allows for faster, more efficient virus propagation and flexibility in choosing influenza target strains. The LMN assay can be performed in as few as 20 hours from start to finish, where traditional MN assays can take days to complete. The luminometer takes only a few minutes to produce a readout, much shorter than staining procedures during MN.

The LMN assay and the HAI assay measure slightly different antibody activities. While the HAI assay quantifies inhibition of hemagglutination, a proxy of virus-cell binding, the LMN assay measures inhibition of cell entry, a proxy of neutralization^10,11^. The LMN assay is more sensitive and has a broader dynamic range compared to the HAI assay (data not shown). For this reason, we standardized LMN NT_75_ to the range of HAI titers by expressing the LMN NT_75_ as ordinal values and setting our “first” titer to the value one titer greater (1:40) than the negative control output (1:20). This allowed for better comparisons across assays. After standardization, we found that LMN NT_75_ and HAI titer are well correlated, suggesting that the LMN assay is a viable alternative to the HAI assay. We did find that the correlation was lower for A/H3N2/Hong Kong/4801/2014 compared to A/H3N2/Singapore/INFIMH-16-0019/2016. These differing correlations can potentially be explained by altered sialic acid binding and hemagglutination properties of the A/H3N2/Hong Kong/4801/2014 HA, highlighting the advantage of using a cell-based LMN assay^13^.

The LMN assay has several limitations. While reverse genetics have been widely used to generate virus, these lab-generated viral strains do have key differences compared to circulating viruses. For example, the viruses we created are on a WSN33 backbone and contain cloned HA and NA segments. With these key differences, there is potential for inconsistencies in antibody binding to these strains compared to the same strains found in nature. However, all HA and NA segments were verified via Sanger sequencing and these methods have been shown to be an acceptable alternative to using primary isolates^13,21^. Additionally, a luminometer is needed to perform this assay, which is an additional expense over the HAI assay. The validation study itself also was subject to limitations. First, our egg-adapted influenza stocks were propagated on MDCK-SIAT1 cells and not chicken eggs. The cellular glycans are different from those in eggs. However, we sequenced our final stocks of virus to ensure that our “egg-adapted” stocks still contained the previously documented egg-adapted genetic changes^7,22,23^. The LMN assay also was able to detect 2-3-fold differences in NT_75_ for egg- and cell-adapted viruses, which suggests further utility for a wide-range of differing influenza targets. Lastly, we are unable to draw conclusions regarding outside factors that may impact protection toward influenza. This study was performed for assay development and validation, with the samples being selected based on discordant HAI titer results to differing H3N2 influenza viruses (data not shown). Because of potential bias, we can’t draw conclusions from these data related to the impact of vaccination or infection history on HAI titer or LMN NT_75_. However, our results provide confidence that this method has the resolution needed to further delve into the impacts of vaccination and infection history on protection.

In summary, the LMN assay was able to produce similar results to the gold-standard HAI assay, as well as detect differences in neutralization response for egg- and cell-adapted influenza strains. Together, these results demonstrate the utility of this assay for studies of immune correlates of protection and highlight key antigenic differences between egg-adapted vaccine strains and their circulating counterparts.

## Acknowledgements

We thank all individuals who participated in this study. This project has been funded in in part with Federal funds from the National Institute of Allergy and Infectious Diseases, National Institutes of Health, Department of Health and Human Services, under Contract No. 75N93021C00015 and R01 AI148371 and from the Centers for Disease Control and Prevention, under U01IP001034.

## References

1. Rolfes MA, Foppa IM, Garg S, et al. Annual estimates of the burden of seasonal influenza in the United States: A tool for strengthening influenza surveillance and preparedness. Influenza Other Respir Viruses. 2018;12(1):132–137. doi:10.1111/irv.12486

2. Jackson ML, Phillips CH, Wellwood S, et al. Burden of medically attended influenza infection and cases averted by vaccination - United States, 2016/17 through 2018/19 influenza seasons. Vaccine. 2022;40(52):7703–7708. doi:10.1016/j.vaccine.2022.11.011

3. Chung JR, Rolfes MA, Flannery B, et al. Effects of Influenza Vaccination in the United States During the 2018-2019 Influenza Season. Clin Infect Dis Off Publ Infect Dis Soc Am. 2020;71(8):e368–e376. doi:10.1093/cid/ciz1244

4. Flannery B, Kondor RJG, Chung JR, et al. Spread of Antigenically Drifted Influenza A(H3N2) Viruses and Vaccine Effectiveness in the United States During the 2018–2019 Season. J Infect Dis. 2020;221(1):8–15. doi:10.1093/infdis/jiz543

5. Tenforde MW, Kondor RJG, Chung JR, et al. Effect of Antigenic Drift on Influenza Vaccine Effectiveness in the United States—2019–2020. Clin Infect Dis. 2021;73(11):e4244–e4250. doi:10.1093/cid/ciaa1884

6. Chung JR, Flannery B, Thompson MG, et al. Seasonal Effectiveness of Live Attenuated and Inactivated Influenza Vaccine. Pediatrics. 2016;137(2):e20153279. doi:10.1542/peds.2015-3279

7. Zost SJ, Parkhouse K, Gumina ME, et al. Contemporary H3N2 influenza viruses have a glycosylation site that alters binding of antibodies elicited by egg-adapted vaccine strains. Proc Natl Acad Sci. 2017;114(47):12578–12583. doi:10.1073/pnas.1712377114

8. Tenforde MW, Patel MM, Lewis NM, et al. Vaccine Effectiveness Against Influenza A(H3N2)– Associated Hospitalized Illness: United States, 2022. Clin Infect Dis. Published online November 3, 2022:ciac869. doi:10.1093/cid/ciac869

9. Levine MZ, Martin ET, Petrie JG, et al. Antibodies Against Egg- and Cell-Grown Influenza A(H3N2) Viruses in Adults Hospitalized During the 2017-2018 Influenza Season. J Infect Dis. 2019;219(12):1904–1912. doi:10.1093/infdis/jiz049

10. Hirst GK. The Quantitative Determination of Influenza Virus and Antibodies by Means of Red Cell Agglutination. J Exp Med. 1942;75(1):49–64. doi:10.1084/jem.75.1.49

11. WHO. Manual for the laboratory diagnosis and virological surveillance of influenza. Published online 2011. https://apps.who.int/iris/handle/10665/44518

12. Sicca F, Martinuzzi D, Montomoli E, Huckriede A. Comparison of influenza-specific neutralizing antibody titers determined using different assay readouts and hemagglutination inhibition titers: good correlation but poor agreement. Vaccine. 2020;38(11):2527–2541. doi:10.1016/j.vaccine.2020.01.088

13. Allen JD, Ross TM. H3N2 influenza viruses in humans: Viral mechanisms, evolution, and evaluation. Hum Vaccines Immunother. 2018;14(8):1840–1847. doi:10.1080/21645515.2018.1462639

14. Okuno Y, Tanaka K, Baba K, Maeda A, Kunita N, Ueda S. Rapid focus reduction neutralization test of influenza A and B viruses in microtiter system. J Clin Microbiol. 1990;28(6):1308–1313. doi:10.1128/jcm.28.6.1308-1313.1990

15. Monto AS, Malosh RE, Evans R, et al. Data resource profile: Household Influenza Vaccine Evaluation (HIVE) Study. Int J Epidemiol. 2019;48(4):1040–1040g. doi:10.1093/ije/dyz086

16. Matrosovich M, Matrosovich T, Carr J, Roberts NA, Klenk HD. Overexpression of the alpha-2,6-sialyltransferase in MDCK cells increases influenza virus sensitivity to neuraminidase inhibitors. J Virol. 2003;77(15):8418–8425. doi:10.1128/jvi.77.15.8418-8425.2003

17. Truelove S, Zhu H, Lessler J, et al. A comparison of hemagglutination inhibition and neutralization assays for characterizing immunity to seasonal influenza A. Influenza Other Respir Viruses. 2016;10(6):518–524. doi:10.1111/irv.12408

18. Trombetta CM, Remarque EJ, Mortier D, Montomoli E. Comparison of hemagglutination inhibition, single radial hemolysis, virus neutralization assays, and ELISA to detect antibody levels against seasonal influenza viruses. Influenza Other Respir Viruses. 2018;12(6):675–686. doi:10.1111/irv.12591

19. Grund S, Adams O, Wählisch S, Schweiger B. Comparison of hemagglutination inhibition assay, an ELISA-based micro-neutralization assay and colorimetric microneutralization assay to detect antibody responses to vaccination against influenza A H1N1 2009 virus. J Virol Methods. 2011;171(2):369–373. doi:10.1016/j.jviromet.2010.11.024

20. Tran V, Moser LA, Poole DS, Mehle A. Highly sensitive real-time in vivo imaging of an influenza reporter virus reveals dynamics of replication and spread. J Virol. 2013;87(24):13321–13329. doi:10.1128/JVI.02381-13

21. Rajaram S, Van Boxmeer J, Leav B, Suphaphiphat P, Iheanacho I, Kistler K. 2556. Retrospective Evaluation of Mismatch From Egg-Based Isolation of Influenza Strains Compared With Cell-Based Isolation and the Possible Implications for Vaccine Effectiveness. Open Forum Infect Dis. 2018;5(suppl_1):S69–S69. doi:10.1093/ofid/ofy209.164

22. Harding A, Heaton N. Efforts to Improve the Seasonal Influenza Vaccine. Vaccines. 2018;6(2):19. doi:10.3390/vaccines6020019

23. Barr IG, Donis RO, Katz JM, et al. Cell culture-derived influenza vaccines in the severe 2017–2018 epidemic season: a step towards improved influenza vaccine effectiveness. Npj Vaccines. 2018;3(1):44. doi:10.1038/s41541-018-0079-z

